# Self-assembling Gn head ferritin nanoparticle vaccine provides full protection from lethal challenge of Dabie Bandavirus in aged ferrets

**DOI:** 10.1101/2023.07.19.549761

**Authors:** Dokyun Kim, Eunha Kim, Semi Kim, Youseung Chung, Sung-Dong Cho, Yunseo Choi, Chih-Jen Lai, Xinghong Dai, Seokmin Kang, Mi-Jeong Kwak, Inho Cha, Ziyi Liu, Younho Choi, Su-Hyung Park, Young Ki Choi, Jae U. Jung

## Abstract

Dabie Bandavirus (DBV), previously known as Severe Fever with Thrombocytopenia Syndrome (SFTS) Virus, induces a characteristic thrombocytopenia with a mortality rate ranging from 12% to as high as 30%. The sero-prevalence of DBV in healthy people is not significantly different among age groups, but clinically diagnosed SFTS patients are older than ∼50 years, suggesting that age is the critical risk factor for SFTS morbidity and mortality. Accordingly, our immune-competent ferret model demonstrates an age (>4 years old)-dependent DBV infection and pathogenesis that fully recapitulates human clinical manifestation. To protect the aged population from DBV-induced SFTS, vaccine should carry robust immunogenicity with high safety profile. Previous studies have shown that glycoproteins Gn/Gc are the most effective antigens for inducing both neutralizing antibody (NAb)- and T cell-mediated immunity and, thereby, protection. Here, we report the development of a protein subunit vaccine with 24-mer self-assembling ferritin (FT) nanoparticle to present DBV Gn head region (GnH) for enhanced immunogenicity. Anion exchange chromatography and size exclusion chromatography readily purified the GnH-FT nanoparticles to homogeneity with structural integrity. Mice immunized with GnH-FT nanoparticles induced robust NAb response and T-cell immunity against DBV Gn. Furthermore, aged ferrets immunized with GnH-FT nanoparticles were fully protected from DBV challenge without SFTS symptoms such as body weight loss, thrombocytopenia, leukopenia, and fatality. This study demonstrates that DBV GnH-FT nanoparticles provide an efficient vaccine efficacy in mouse and aged ferret models and should be an outstanding vaccine candidate targeted for the aged population against fatal DBV infection.

**Importance:** Dabie Bandavirus (DBV) is an emerging tick-borne virus that causes Severe Fever with Thrombocytopenia Syndrome (SFTS) in infected patients. Human SFTS symptoms progress from fever, fatigue, and muscle pain to the depletion of white blood cells and platelets with fatality rates up to 30%. The recent spread of its vector tick to over 20 states in the United States increases the potential for outbreaks of the SFTS beyond the East Asia. Thus, the development of vaccine to control this rapidly emerging virus is a high priority.

In this study, we applied self-assembling ferritin (FT) nanoparticle to enhance the immunogenicity of viral Gn head domain as a vaccine target. Mice immunized with the GnH-FT nanoparticle vaccine induced potent antibody responses and cellular immunity. Immunized aged-ferrets were fully protected from the lethal challenge of DBV. Our study describes the GnH-FT nanoparticle vaccine candidate that provides protective immunity against the emerging DBV infection.

## Introduction

Dabie Bandavirus (DBV), previously known as Severe Fever with Thrombocytopenia Syndrome (SFTS) Virus (SFTSV), is an emerging Bunyavirus responsible for causing SFTS in infected individuals (1). Since its initial identification in 2009 (2), the virus has established endemic infections in China, South Korea, and Japan, and has recently expanded to Southeast Asia (1, 3, 4). Prognoses of human DBV infection begin with flu-like symptoms, such as fever, fatigue, myalgia, and progress to hemorrhagic manifestations, including leukopenia, thrombocytopenia, and multiorgan failure, with fatality rates ranging from 12% to 30% (1, 5, 6). The severity of SFTS and its outcomes exhibit a clear age-dependence, as the vast majority of fatal cases and hospitalizations occur in individuals aged 50 or older (7, 8). However, no licensed vaccine or therapy against DBV is currently available. Consequently, the World Health Organization (WHO) and the United States National Institute of Allergy and Infectious Diseases (NIAID) have recently designated DBV as one of their priority pathogens (9) and Category C agents (10), respectively, representing emerging pathogens with outbreak potential. This designation aims to stimulate research interest in the development of vaccines and therapies for DBV.

**Fig 1.**
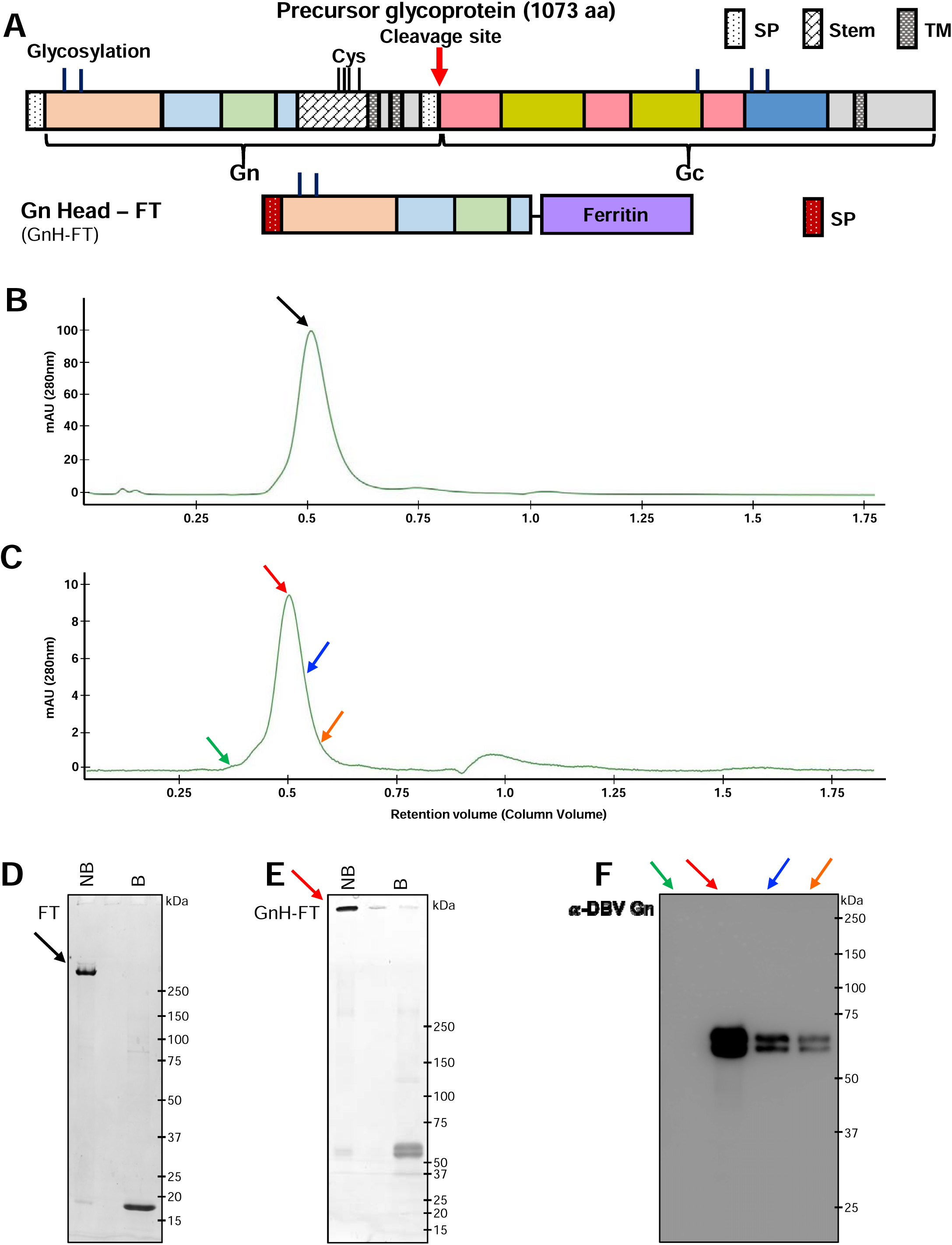
Molecular design and biochemical and antigenic characterization of ferritin nanoparticles (FT) and DBV Gn Head-ferritin (GnH-FT) nanoparticles. (A) Schematic representation of GnH-FT based on previously solved structures and domains of DBV Gn and Gc. The construct was transfected to HEK293T cells to collect the cell supernatant at 72 hours after transfection for purification. (SP: signal peptide, TM: transmembrane domain) (B, C) Size exclusion chromatograms to purify FT (B) and GnH-FT (C) using Superdex 200. Increase 10/300 GL and Superose6 Increase 10/300 GL columns, respectively, on Bio-Rad NGC chromatography system. Fractions corresponding to the colored arrows were separately collected for further analyses. (D, E) Fractions from size exclusion chromatograms of FT and GnH-FT were further analyzed by gradient (7% - 20%) SDS-PAGE and Coomassie Brilliant blue staining. Fractions corresponding to the black and red arrows from FT and GnH-FT purifications were loaded to SDS-PAGE gel without boiling (“NB”) and with boiling (“B”) to characterize head-mediated disassembly of the 24-mer nanoparticle. Intact FT nanoparticle and GnH-FT nanoparticle each has expected molecular weight of approximately 432 kDa and 1,560 kDa. Dissembled FT and GnH-FT monomer each has expected molecular weight of 18 kDa and 65 kDa. (F) Western-blot analysis of different fractions collected from GnH-FT size exclusion chromatogram. In house-generated mouse monoclonal antibody recognizing DBV Gn Head region was used to detect GnH-FT subunit monomers.

DBV belongs to the *Bandavirus* genus within the *Phenuiviridae* family of the *Bunyavirales* order and carries a single-stranded, negative-sense genome divided into three segments: L, M, and S. The M segment encodes a precursor glycoprotein Gn/Gc, which is subsequently processed into Gn and Gc by host proteases (11). The Gn and Gc proteins assemble highly ordered capsomers as dimers on the viral surface. Gn is responsible for viral attachment to host cells through receptor binding, enabling Gc to mediate membrane fusion (12). Previous studies have identified DBV-neutralizing antibody epitopes on the head region of Gn (13–15), strongly suggesting the potential of DBV Gn as a vaccine candidate. Furthermore, our prior efforts to develop a DBV DNA vaccine demonstrated protection against lethal DBV challenge and the most potent induction of immunity when immunized with the M segment, which encodes the glycoproteins (16).

There has been limited success in developing vaccines against DBV due to the absence of an immunocompetent animal model that replicates clinical symptoms from human DBV infection and subsequent SFTS pathogenesis. In particular, the failure to reproduce age-dependent disease progression and clinical outcomes has been a significant obstacle in developing a vaccine to protect the most vulnerable group – the elderly population. We have recently presented an immunocompetent ferret model that highly replicates human DBV infection and clinical symptoms of SFTS. Aged ferrets (4 years or older) fully recapitulates SFTS disease pathologies, characterized by fever, thrombocytopenia, leukopenia, and viremia in blood and organs, resulting in 93% fatality rate (17). Aged ferret model have been applied to DBV vaccine development using a live-attenuated vaccine with a mutation in its major virulence factor (non-structural protein, NSs) (18) and a DNA vaccine encoding the M segment to express Gn and Gc (16). Immunized aged ferrets developed strong humoral and cellular immunity and were fully protected from lethal DBV challenge.

The most vulnerable group to DBV infection and fatal SFTS is the elderly population, which experiences suboptimal induction of immunity and a high risk of vaccine-related adverse effects (19). Therefore, a vaccine development approach with an excellent safety profile and immunogenicity is required to effectively protect the elderly population against DBV infections. Protein subunit vaccines of viral antigens containing neutralization epitopes have been suggested as safe vaccine candidates; however, host immune system often fails to effectively react against soluble antigens due to their small sizes (20). Fortunately, recent advances nanotechnology and molecular biology have transcended previous limitations by employing nanoparticle engineering as a toolkit for vaccine development (21). The immunogenicity of nanoparticle-engineered vaccines has surpassed that of traditional protein subunit vaccines. Moreover, recent research has elucidated the immunological mechanisms behind the enhanced immunogenicity: higher activation and formation of germinal centers (22), improved antigen transport to draining lymph nodes (23), and antigen presentation by follicular dendritic cells and helper T cells (24). Among the naturally derived and artificially designed nanoparticles, ferritin is the most extensively studied and applied nanoparticle. Found across life’s kingdoms, ferritin possesses a conserved function in storing excess iron (Fe^2+^) inside the nanoparticle to quench the Fenton reaction, which generates reactive oxygen species that cause cellular damage. More importantly, ferritin forms a higher-order homopolymer structure of the self-assembly of 24 ferritin monomer subunits, facilitating expression and purification for application in biotechnology. Its molecular amenability via fusion peptides from recombinant DNA constructs has enabled further application of the self-assembling nanoparticle to vaccine development (25).

One of the most commonly applied ferritins is hybrid ferritin, engineered by fusing the *Helicobacter pylori* ferritin backbone with NH_2_-terminal tail from bullfrog (*Rana catesbeiana*) ferritin lower subunit. The NH_2_-terminal tail forms radial projections on the threefold-axis points of the self-assembled nanoparticle, efficiently presenting viral immunogens fused in the recombinant DNA construct to provide stronger protective immunity at significantly lower doses than soluble antigens (26, 27). Due to the low amino acid sequence similarity of *H. pylori* and bullfrog ferritin to human ferritin, the hybrid ferritin has minimal risk of vaccine-related adverse effects or autoimmunity from antigen mimicry. Consequently, the hybrid nanoparticle has been widely applied as an effective immunogen carrier platform in diverse vaccine developments, including MERS-CoV (28), SARS-CoV-2 (25), Influenza (27), and Epstein-Barr Virus (26). These indicate the utility of ferritin nanoparticles as an outstanding vaccine carrier platform to induce strong protective immunity against infectious agents.

In this paper, we demonstrate the immunogenicity of the self-assembling DBV Gn Head (GnH)-ferritin (GnH-FT) nanoparticle as an effective DBV vaccine candidate. We purified GnH-FT nanoparticles from transfected HEK293T cells, characterized their biochemical, antigenic and structural profiles, and immunized mice and aged ferrets to investigate the induction of humoral and cellular immunity. Furthermore, we challenged vaccinated aged ferrets with a lethal dose of DBV and observed protective immunity against DBV. These data suggest the DBV GnH-FT nanoparticle as a promising vaccine candidate, providing protective immunity against DBV infection and subsequent SFTS pathogenesis.

## Results

### Molecular design, purification, and characterization of DBV GnH-ferritin nanoparticles

A majority of DBV-neutralizing antibodies from human convalescent sera target the head region of Gn (13, 14), which mediates viral attachment to host cells. Furthermore, our recent study of DBV DNA vaccine has identified the viral glycoproteins as the most immunogenic antigen among the viral proteins. The DBV Gn head (GnH) gene was first human-codon-optimized and fused to the IL-2 signal peptide at the N-terminus and the *H. pylori*-bullfrog hybrid ferritin at the C-terminus to generate GnH-ferritin (GnH-FT) (Fig. 1A). HEK293T cells were transfected with GnH-FT expression plasmid or the hybrid ferritin (FT) with the signal peptide, and the cell supernatants were collected to purify the GnH-FT and FT nanoparticles by anion exchange chromatography and size exclusion chromatography.

The purified FT (Fig. 1B) and GnH-FT (Fig. 1C) nanoparticles were homogenous, as demonstrated by size exclusion chromatography with specific columns exhibiting maximal separating resolution at several hundred kilodaltons and a few megadaltons, respectively. Additionally, the chromatograms of GnH-FT and FT nanoparticles showed peaks at fractions corresponding to the expected molecular weight as 24-mer nanoparticles. We further tested if the purified FT and GnH-FT nanoparticles retained the 24-mer nanoparticle structures by loading the purified fractions onto SDS-PAGE without boiling (“NB” for not boiled) or with boiling (“B” for boiled). The purified nanoparticles retained the higher-order structure without boiling but disassembled into subunit monomers upon boiling (Fig. 1D, 1E). As a result, bands appeared at the expected molecular weights of monomers calculated based on previously solved GnH structures (15). Immunoblotting analysis of different fractions (marked with colored arrows) of size exclusion chromatography with anti-DBV Gn antibody also confirmed the presence of GnH-ferritin with intensities matching the peak heights of size exclusion chromatogram (Fig. 1F). These results indicate that the purified GnH-FT nanoparticles retain the higher-order structure from self-assembly.

### Purified GnH-FT nanoparticle retains the higher-order structure and presents GnH on its surface

According to computer-assisted modeling based on previous reports using ferritin nanoparticles as carrier platforms, the GnH antigen was expected to radially project from three-fold axis points of the nanoparticle (Fig. 2A, 2B). Negative staining transmission electron microscopy (EM) and cryo-EM of FT nanoparticles demonstrated a homogenously smooth circular surface with an average diameter of 9.5nm (Fig. 2C, Fig. S1A), while those of GnH-FT nanoparticles showed clearly visible protrusions from the ferritin core with an average diameter of 14.7nm (Fig. 2D and Fig. S1B). These protrusions appeared as an extra layer of white, smeared halo surrounding the ferritin core in the cryo-EM 2D class averages of GnH-FT nanoparticles (Fig. 2F) compared to those of the FT nanoparticles (Fig. 2E). These data indicated that the GnH was flexible on the surface of the ferritin particle, and its presence did not affect the assembly of the ferritin particle.

**Fig 2.**
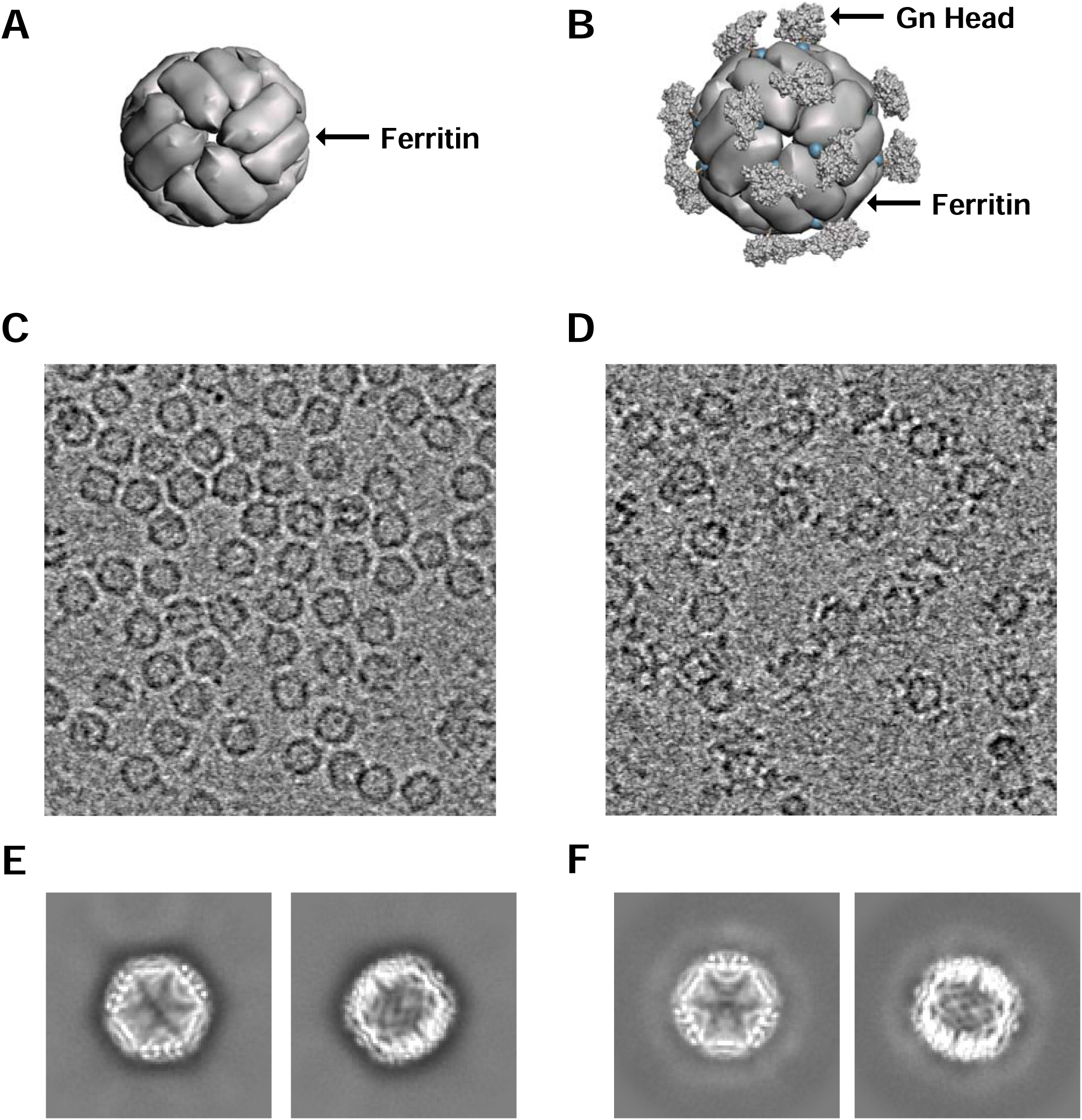
Prediction and observed structures of FT and GnH-FT nanoparticles. (A, B) Computer-assisted 3D model of the FT nanoparticles (A) and GnH-FT nanoparticles (B) based on previously solved structures of DBV Gn (PDB: 5Y11) and FT nanoparticles (PDB: 3EGM). (C, D) Cryo-electron microscopy (cryo-EM) of FT (C) and GnH-FT (D) nanoparticles. (E, F) Representative 2D class averages of FT (E) and GnH-FT (F). The white halo surrounding the FT nanoparticle core in (F) is attributable to the highly flexible linker and DBV Gn head domain.

### Immunization with GnH-FT nanoparticle induces humoral immunity and cellular immunity *in vivo*

To enhance the relatively weaker immunogenicity of protein subunit vaccines, a diverse selection of adjuvants is administered in combination with protein vaccine candidate. One of the safest among the variety of adjuvants is MF59, an oil-in-water emulsion adjuvant used in adjuvanted flu vaccines (29–32). We combined the veterinary equivalent MF59, AddaVax, with FT or GnH-FT nanoparticles for immunization. 8-10-week-old BALB/c mice (n=6 per antigen) were immunized with a total of 3 doses at 3-week intervals of 3.3μg of FT nanoparticle – equimolar to 10μg of GnH-FT nanoparticle – or 1μg, 5μg, or 10μg of GnH-FT nanoparticle via the intramuscular route. Blood was drawn prior to immunization (week 0) and every week starting 2 weeks after the priming immunization. Total IgG antibody against GnH soluble protein reached a maximal level at 2 weeks after the 2^nd^ dose of immunization (first booster) and did not further increase upon the 3^rd^ dose (second booster) among all three doses of GnH-FT nanoparticle immunization (Fig. 3B). Interestingly, there was no statistically significant difference in the induction of total anti-GnH IgG levels across different doses of GnH-FT nanoparticle, indicating that 1μg GnH-FT dose was sufficient for inducing strong antibody response.

**Fig 3.**
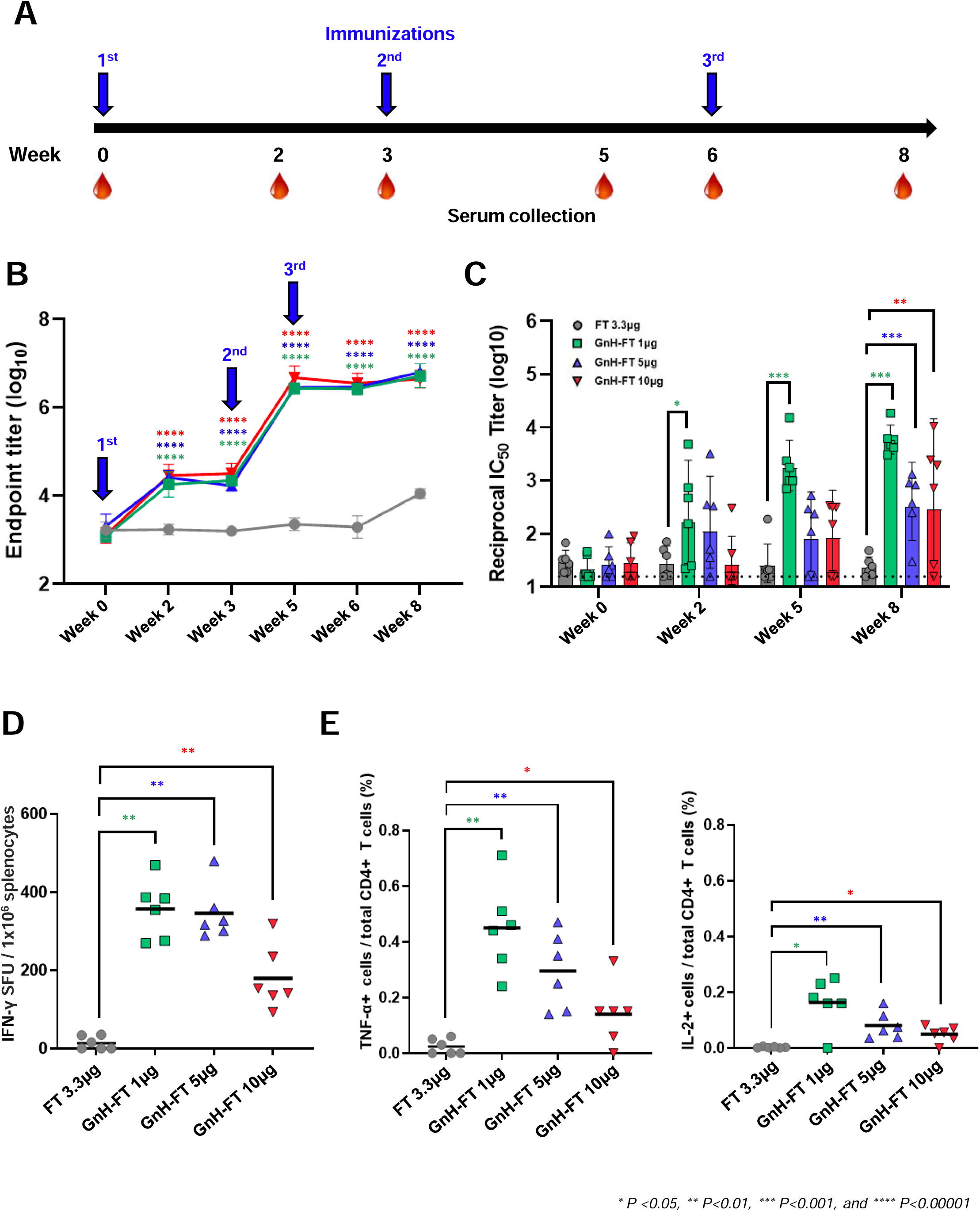
Immunization with GnH-FT elicits humoral and cellular immunity *in vivo*. (A) Timeline for mouse immunization and blood collection. Six BALB/c mice per antigen group were intramuscularly immunized at hind leg with 3.3μg of FT or 1μg, 5μg, or 10μg of GnH-FT. 3.3μg of FT is equimolar to 10μg of GnH-FT. (B) Reciprocal IgG titer measured by ELISA using purified GnH-10His protein coated. Sera from blood samples collected at weeks 0, 2, 3, 5, 6, and 8 were used to quantify total IgG recognizing DBV Gn Head. Endpoint titer values are presented in log_10_ values. The asterisks represent statistical significance of endpoint titer between mice immunized with GnH-FT and FT evaluated with one-way ANOVA with Dunnett multiple comparison test. (C) Neutralizing antibody titer measured by luciferase assay using rVSV-DBV G carrying luciferase gene. Sera from blood samples collected at weeks 0, 2, 5, and 8 were used to quantify reciprocal IC_50_ titer to represent induction of neutralizing antibody upon immunization with different antigens. One-way ANOVA with Dunnett multiple comparison test was performed. (D) ELISpot assays were performed to detect DBV Gn Head-specific T cells secreting IFN-γ by *ex vivo* stimulation of whole splenocytes with pool of overlapping peptides (OLP). Data represents the number of spot-forming units (SFUs) per 1 million splenocytes. One-way ANOVA with Dunnett multiple comparison test was used to evaluate statistical significance. (E) Intracellular cytokine staining assay was performed to test for activity of cellular immunity induced upon immunization with the antigens. Splenocytes were stimulated *ex vivo* as for ELISpot assay, treated with protein transport inhibitor, then stained for anti-TNF-α antibodies or IL-2 antibodies. Two-tailed, unpaired t-test was performed to evaluate statistical significance. *P < 0.05, **P < 0.01, ***P < 0.001 and ****P < 0.00001.

We performed a neutralization assay using replication-defective recombinant Vesicular Stomatitis Virus (rVSV) carrying DBV glycoproteins, Gn and Gc, and luciferase reporter gene (rVSV-DBV G). Unlike total anti-GnH antibody response that reached a maximal level after the 2^nd^ immunization, neutralizing antibody (NAb) titer continuously increased over the 3 immunizations with GnH-FT nanoparticle (Fig. 3C). Immunization with 1μg GnH-FT nanoparticle elicited the most robust neutralizing antibody response against DBV, followed by 5μg and 10μg. Consistently with the total IgG, there was no significant induction of NAb against DBV upon immunization with FT nanoparticle alone (Fig. 3C). These data suggest that 1μg GnH-FT may be an optimal dose for 3-dose regimen to elicit strong NAb response against DBV infection.

Although NAb titer is a critical marker of vaccine efficacy, many studies have shown the importance of cellular immunity for antiviral immunity (33, 34). To perform IFN-γ ELISpot, spleens from immunized mice at week 8 (two weeks after the 3^rd^ immunization) were harvested, *ex vivo* stimulated with a pool of overlapping peptides (OLPs) spanning GnH, and subsequently subjected to IFN-γ ELISpot. This showed that IFN-γ secretion was induced across all doses of GnH-FT nanoparticle immunization, whereas immunization with 1μg GnH-FT nanoparticle elicited the most robust IFN-γ secretion (Fig. 3D). In addition, GnH-FT nanoparticle immunization successfully induced TNF-α and IL-2 production from OLP-stimulated CD4+ T cells, whereas FT-nanoparticle immunization did not induce those cytokine productions (Fig. 3E). Finally, there was no significant production of TNF-α and IL-2 from OLP-stimulated CD8+ T cells (data not shown). These showed that the maximal induction of NAb, IFN-γ, TNF-α and IL-2 were observed from 1μg dose of GnH-FT nanoparticle immunization. The data corresponds to previous reports of robust activation of protective immunity from immunization with nanoparticle at significantly lower doses. Collectively, these results demonstrate that the immunization of DBV GnH-FT nanoparticle effectively elicits both NAb production and T-cell response in mice.

### Aged ferret immunized with GnH-FT nanoparticle form antibody responses against DBV

Naïve 4-year-old ferrets (n=12 ferrets per antigen) were vaccinated via intramuscular injection with FT or GnH-FT nanoparticles with AddaVax adjuvant for a total of 3 immunizations at 2-week intervals. Blood of immunized ferrets was collected on the day of immunization to characterize antibody responses (Fig. 4A). While total anti-GnH IgG titers were dramatically increased after the 1^st^ or 2^nd^ vaccination, they were further escalated after the 3^rd^ vaccination, suggesting that the vaccination protocol of total 3 doses maximizes the antibody response in aged ferret. We did not observe significant increase of IgG level upon booster immunizations and it is attributable to saturation of the assay (Fig. 4B). Consistently, serum NAb titers against DBV CB1/2014 strain were continuously increased following the priming vaccination and subsequent booster vaccinations (Fig. 4C). On the contrary, neither anti-GnH antibody nor anti-DBV NAb was detected from control ferrets immunized with FT nanoparticles. These data indicate that the GnH-FT nanoparticle effectively induces antibody responses against DBV in aged ferret model.

**Fig 4.**
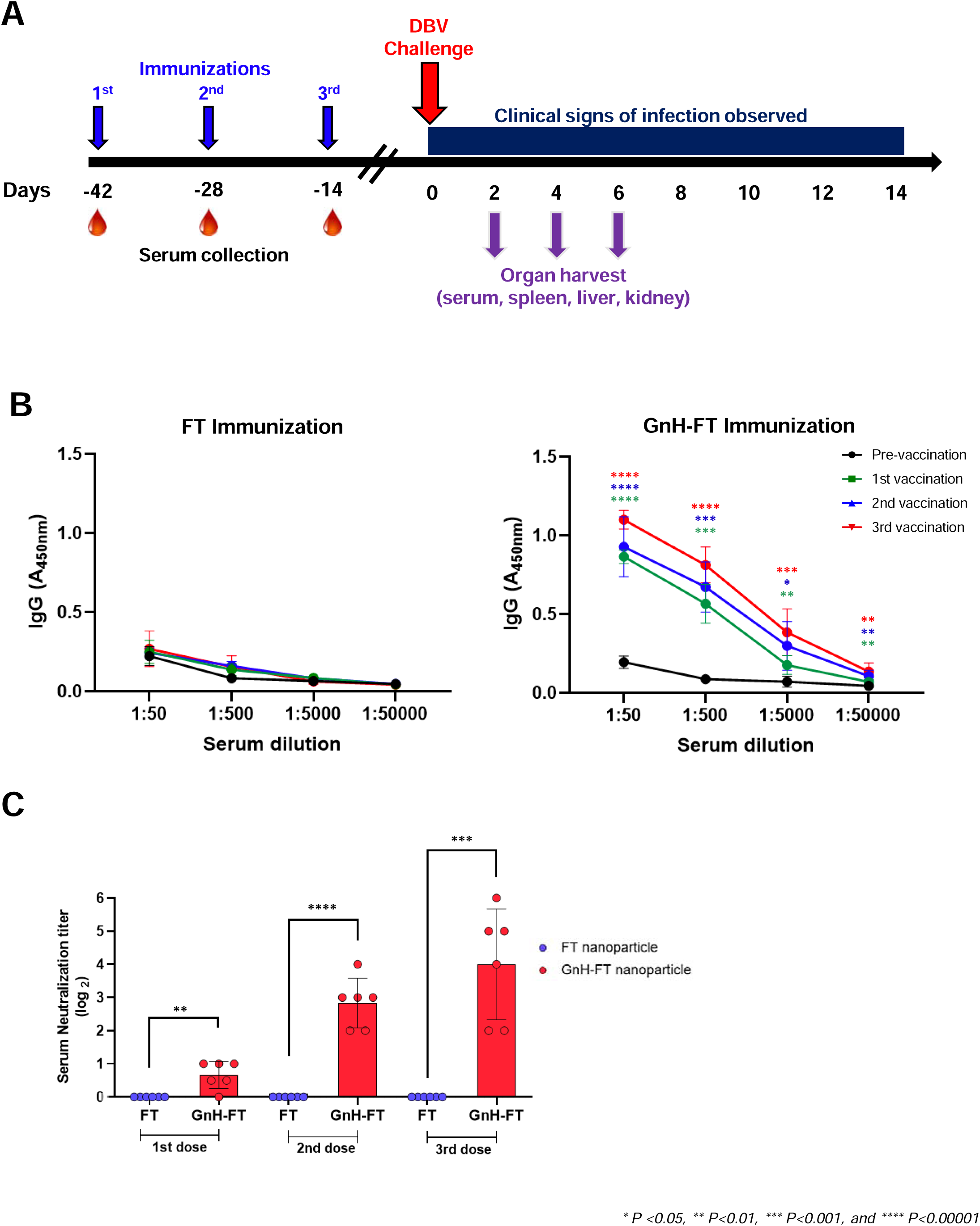
Aged ferrets form humoral immunity upon immunization with GnH-FT. (A) Timeline for aged ferret immunization, lethal DBV challenge, and organ harvest. Based on the effective induction of humoral and cellular immunity in mouse model, aged ferrets were immunized with total three doses of 15μg GnH-FT or FT. Each antigen group had 12 ferrets. Aged ferrets were challenged with 10^7.6^ TCID_50_/mL of DBV for infection with lethal dose at 2 weeks after the last booster immunization, then monitored for clinical symptoms of SFTS. At 2, 4, and 6 days post the challenge, three ferrets per antigen group were sacrificed to harvest serum, spleen, liver and kidney for organ virus titration to test for acceleration in viral clearance. (B) Reciprocal IgG titer measured by ELISA using blood samples collected at days −42, −28, −14 and 0 to characterize humoral immunity induced by immunization with FT or GnH-FT. Optical density (OD) was measured with a spectrometer (VarioSkan, Thermo) at detection wavelength of 450nm.The asterisks represent statistical significance of OD measurements from ferrets immunized with GnH-FT to ferrets immunized with FT evaluated with one-way ANOVA with Dunnett multiple comparison test. (C) Neutralizing antibody response to DBV elicited by immunization with FT or GnH-FT was characterized as FRNT_50_ from blood samples collected at days −28, −14, and 0 to represent time points of 2 weeks after the 1^st^, 2^nd^, and 3^rd^ vaccination. The asterisks represent statistical significance of FRNT_50_ from ferrets immunized with GnH-FT to ferrets immunized with FT evaluated with one-way ANOVA with Dunnett multiple comparison test. *P < 0.05, **P < 0.01, ***P < 0.001 and ****P < 0.00001.

### Immunization with GnH-FT nanoparticle provides full protection against lethal DBV challenge in aged ferrets

To evaluate the protective efficacy of GnH-FT vaccines, FT or GnH-FT vaccinated aged ferrets were intramuscularly challenged with a lethal dose of DBV CB1/2014 strain (10^7.6^ TCID50) in 2 weeks after the third vaccination, and monitored for clinical signs of infection, viral titers and platelet counts in the blood, body weight, body temperature, and survival rate for the following 14 days, with evaluations every other day. Blood was collected every other day to measure platelet and white blood cell counts to observe thrombocytopenia and leukopenia (Fig. 4A).

Strikingly, all ferrets immunized with GnH-FT nanoparticles were fully protected from lethal DBV challenge, while ferrets immunized with FT nanoparticles suffered significant body weight loss up to 20% and succumbed to death (Fig. 5A and 5B). Aged ferrets immunized with GnH-FT nanoparticles showed minimal increases in body temperature, while those immunized with FT nanoparticles experienced severe fever (Fig. 5C). Platelet and white blood cell counts were also measured from the blood samples to test for the characteristic symptoms of thrombocytopenia and leukopenia. Consistent with body weight, temperature, and survival, aged ferrets immunized with GnH-FT nanoparticles showed little or no significant reduction in platelet and white blood cell counts (Fig. 5D). In contrast, aged ferrets immunized with FT nanoparticles demonstrated dramatic reductions in both platelet and white blood cell counts before the fatal outcome (Fig. 5E).

**Fig 5.**
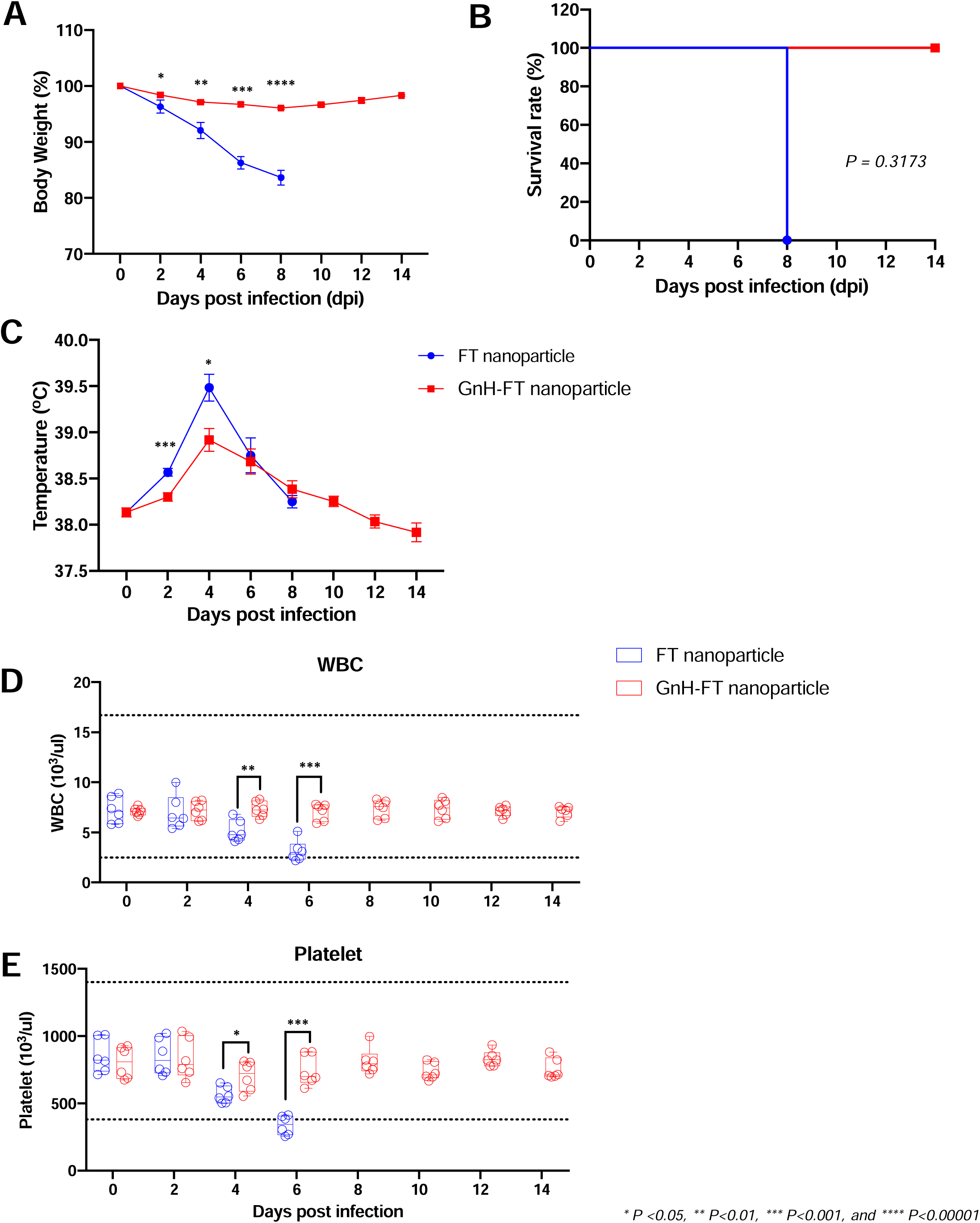
Protective immunity elicited by GnH-FT nanoparticle immunization against lethal DBV challenge in aged ferrets. (A, B, C) Body weight (A), survival curve (B) and temperature (C) of the aged ferrets were observed for 14 days from the lethal challenge. Body weight and temperature are presented as mean ± SEM and statistical significance was analyzed by one-way ANOVA with Dunnett multiple comparison test. Statistical significance of survival across the antigens was analyzed with 2-tailed Mantel-Cox method. (D, E) White blood cell (WBC, D) and platelet (E) counts were measured from blood samples collected for 14 days after the lethal challenge. Data are presented as box plots with the upper (75%) and lower (25%) quartiles, the horizontal line (median) and whiskers (maximum and minimum). Statistical significance across the antigens was evaluated by a two-tailed unpaired t-test. *P < 0.05, **P < 0.01, ***P < 0.001 and ****P < 0.00001.

To assess viral burden in multiple organs upon lethal DBV challenge, we sacrificed 3 ferrets at days 2, 4, and 6 post-infection and harvested serum, spleen, liver, and kidney. Viral titers were measured as RNA copy numbers using real-time PCR. Aged ferrets immunized with FT-nanoparticle demonstrated significant viremia in liver and kidney upon lethal DBV challenge and succumbed to the viral challenge (Fig. 6C and 6D). However, aged ferrets immunized with GnH-FT nanoparticles rapidly cleared the challenging virus so that the viral titer quickly decreased to the limit of detection or lower (Fig. 6C and 6D). These data indicate that immunization with GnH-FT nanoparticles provides complete protection against SFTS pathogenesis upon lethal DBV challenge and promotes viral clearance in aged ferrets.

**Fig 6.**
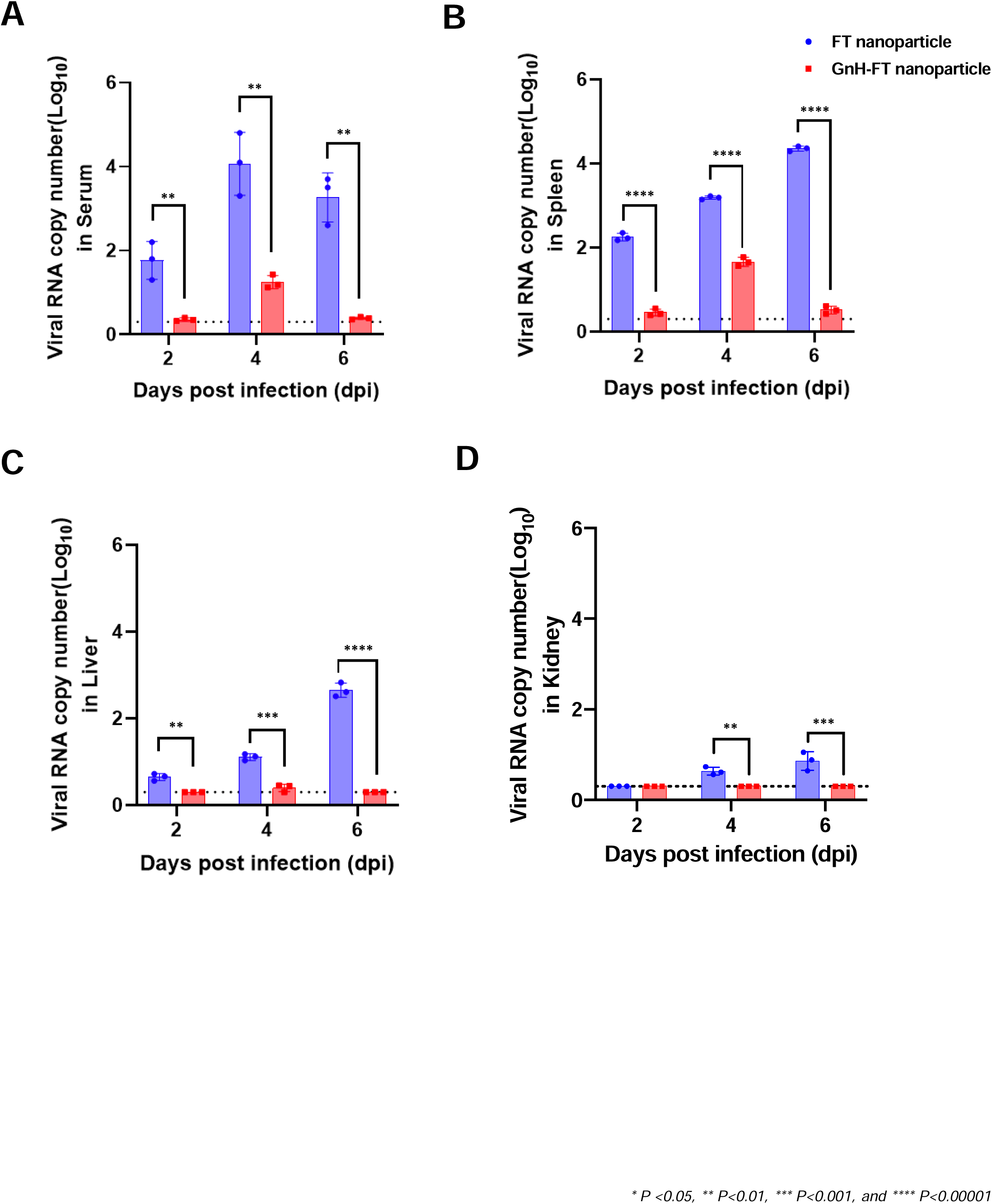
Viral titration from serum, spleen, liver and kidney. Organs were harvested at days 2, 4, and 6 from three animals at each time point. DBV titer values from serum (A), spleen (B), liver (C), and kidney (D) were measured with real-time PCR. Data are shown as mean ± SEM. The asterisks indicate statistical significance compared between ferrets immunized with FT nanoparticle and GnH-FT nanoparticle from the two-tailed, unpaired t-test. *P < 0.05, **P < 0.01, ***P < 0.001 and ****P < 0.00001.

## Discussion

DBV, previously named SFTSV, is an emerging pathogen causing fatal SFTS in infected patients. Since its original discovery in China, it has established endemic infection in South Korea, Japan, and China, and spread to Southeast Asian countries (3, 4, 8). A clear age-dependence in pathogenesis from human DBV infection is evident, as the majority of hospitalization cases and almost all fatal infections occur in age groups of 50 or above (7). The vector tick – *Haemaphysalis longicornis* – used to have a relatively confined habitat in East Asia. However, its parthenogenetic reproduction has enabled recent rapid spread to other continents, including Australia and North America (35, 36). Combined with the spread of the tick, there have been growing concerns of a DBV outbreak beyond East Asia (1, 7, 36, 37). Here, we demonstrate the immunogenicity of the self-assembling GnH-FT nanoparticle as an effective DBV vaccine candidate. Mice immunized with the FT nanoparticle vaccine induced potent antibody responses and cellular immunity. Immunized aged ferrets were fully protected from the lethal infection of DBV. Our results strongly demonstrate the DBV GnH-FT nanoparticle as a promising vaccine candidate, providing protective immunity against DBV infection and subsequent SFTS pathogenesis.

Among DBV viral proteins, Gn and Gc glycoproteins are initially translated as a precursor glycoprotein from the M segment of the viral genome and subsequently processed into separate proteins by host protease (11). Structural analyses of closely related Bunyaviruses – Heartland Bandavirus (HRTV) and Rift Valley Fever Virus (RVFV) – have identified that Gn and Gc form heterodimers and create higher-order structures on the viral surface (38, 39). Gn and Gc contribute to viral infection with Gn attaching to the host cell membrane and Gc mediating membrane fusion for endocytosis (12, 40). Our previous efforts in developing a DNA vaccine for DBV showed its effectiveness in protecting aged ferrets against fatal DBV infection and eliciting the most robust immune response when immunized with the M segment encoding the glycoproteins (16). Furthermore, all neutralizing antibodies reported to date from convalescent human sera have mapped their epitopes on the head region of Gn (13–15), strongly suggesting its potential as a prime target for vaccine development.

Previous studies have demonstrated higher immunogenicity at lower doses of nanoparticle vaccines compared to conventional protein vaccines while retaining the advantage of reduced reactogenicity (26, 27). To develop a safe yet sufficiently immunogenic vaccine for aged population with weakened immunity, we formulated GnH-FT nanoparticle with an adjuvant with an established safety profile in the elderly population. Purified GnH-FT nanoparticles showed their integrity in presenting the DBV GnH on the carrier nanoparticle. Mice and aged ferrets immunized with GnH-FT nanoparticle robustly induced DBV-recognizing IgG and NAb. While Nab titers increased over booster immunizations, the highest NAb titers were observed from the immunization with 1μg of GnH-FT nanoparticle. Mice immunized with 1μg GnH-FT nanoparticle also showed the strongest T cell response. This corresponds to the previous findings that immunizations with low doses of ferritin-fused nanoparticle provide protective immunity against Influenza virus (0.22μg) (27), Epstein-Barr virus (0.5μg) (26) and SARS-CoV-2 vaccine (15μg) (25). These studies further support the strong immunogenicity of ferritin nanoparticle at low doses as vaccine carrier. From our mouse *in vivo* studies, immunization with 1μg GnH-FT provided the most robust induction of NAbs and T-cell responses. Because dose is closely related to vaccine-related adverse effects, additional studies are needed to optimize GnH-FT immunization protocols to enhance vaccine-mediated immunity while minimizing reactogenicity.

From our aged ferret *in vivo* studies, booster immunizations with GnH-FT nanoparticle induced a significant increase in serum neutralization titer. However, there was only a marginal increase in the total IgG recognizing DBV Gn. This could be due to a potential saturation of ELISA assay performed to capture the total IgG and subsequent limitation in reporting the actual titer.

There are six genotypes of DBV (A to F) reported to date from South Korea, Japan, and China (7). Geographical locations with endemic DBV infection display varying abundances of these genotypes and, therefore, varying fatality rate. The GnH is encoded by the M segment of the DBV genome, which exhibits lower than 10% viral nucleotide variation (41, 42) and 6% amino acid variation (43), although the numbers vary across genotypes. Another study reported broad cross-reactivity of the HB29 strain of DBV (used in this study) with both autologous and heterologous genotypes of DBV (16, 44). Our previous DBV vaccine development also reported cross-protection of live-attenuated vaccines using HB29 (16, 45). This suggests that HB29 may serve as an optimal standard strain for DBV vaccine development. Protective immunity can be expanded even further for broader protection by exchanging the GnH on the FT nanoparticle with the GnH of other genotypes as “plug and play” platform. Future studies on immunity against other genotypes will provide insights into the potential for broad protection by GnH-FT immunization.

FT nanoparticles have been widely applied in biotechnology due to its well understood assembly process, wide application as carrier, and thermal and chemical stability. These advantages facilitate manufacture, storage, and transportation of FT nanoparticle vaccines. Further characterizations of the maintenance of GnH-FT nanoparticle stability will streamline logistics aspects of the vaccine candidate and contribute to developing an effective and accessible vaccine against DBV.

In conclusion, we designed and purified DBV GnH-FT nanoparticle presenting GnH while retaining the structural integrity of nanoparticles and antigenicity of GnH. Furthermore, we evaluated its immunological efficacy as a vaccine candidate in mouse and aged ferret models. Immunized mice with GnH-FT nanoparticle induced strong humoral immunity and cellular immunity. Immunized aged ferrets showed not only effective induction of total IgG antibody and NAb but also full protection from SFTS symptoms and fatality upon lethal DBV challenge. Although previous reports have shown strong efficacies of live attenuated virus vaccines (45) and DNA vaccine (16), these approaches still have safety concerns, especially in elderly population. Our protein subunit FT nanoparticle vaccine that has prospective safety profile demonstrates outstanding protection efficacy in mouse and aged ferret models. This suggests the GnH-FT nanoparticle as a potential safe vaccine targeted for the elderly population that is a prime target of DBV infection and subsequent SFTS.

## Material and Methods

### Expression and purification of the nanoparticles

Expression vectors to purify the nanoparticles were prepared as previously described in our earlier publication (25). Dabie Bandavirus (DBV) glycoprotein Gn gene (GenBank NC_018138.1) was codon-optimized for human codon usage (Genscript) and cloned into the expression vector. At 70% confluency, HEK293T cells (ATCC) cells had their media changed to FreeStyle 293 medium (Gibco) and transfected with the plasmids. Supernatants were concentrated with 100kDa or 500kDa MWCO filters on Labscale TFF (Sigma). Concentrated supernatants were flowed into anion Resource Q column (Cytiva) for anion exchange chromatography on NGC FPLC (Bio-Rad) running with pH8.0, 20mM Tris-Cl and a gradient increase from 0M to 1M NaCl at 3.0ml/min. Fractions from NaCl concentration of 200mM to 500mM were collected and further purified by size exclusion chromatography. NGC FPLC equipped with Superdex 200 Increase 10/300 GL (for FT nanoparticle) and Superose 6 Increase 10/300 GL (for GnH-FT nanoparticle) columns (Cytiva) were used with PBS at 0.1ml/min. Collected fractions were analyzed by loading onto SDS-PAGE with or without boiling for 10min at 95°C and stained with Coomassie Brilliant blue. Western blot of the fractions was performed using in house-generated mouse monoclonal antibody against DBV Gn.

To purify DBV GnH-10His protein, HEK293T cells were transfected with the mammalian expression vector under the same condition as for nanoparticle purification. Supernatant was flowed into HisTrap HP column (Cytiva) using NGC at flow rate of 5ml/min, and eluted by gradient increase of imidazole from 0mM to 500mM in 150mM NaCl and 20mM Tris-Cl, pH8.0. Fractions were tested for yield and purity by SDS-PAGE and stored at −80°C in 10% glycerol.

### Computer-assisted 3D modeling of FT and GnH-FT nanoparticles

Hypothetical structures of FT nanoparticle and GnH-FT nanoparticles were designed based on previously solved DBV Gn (PDB: 5Y11) and *H. pylori*-bullfrog hybrid ferritin (PDB: 3EGM) with Chimera (University of California San Francisco), PyMol (Schroedinger), and Meshmixer (Autodesk). The model was revised to take account for the size ratio of FT nanoparticles and GnH-FT nanoparticles.

### Transmission electron microscopy and cryo-EM analysis of FT and GnH-FT nanoparticles

For negative staining transmission electron microscopy (EM), carbon-coated grids were rendered hydrophilic by glow-discharge and applied with a drop of purified nanoparticles in DPBS. After absorption for 1min, excess sample was blotted away, and the grids were stained with 1% (w/v) uranyl acetate. After drying, the grids were imaged on a Talos F200X G2 microscope at 200kV.

To prepare cryo-EM grid, an aliquot of 3.5µL purified nanoparticles at ∼1mg/mL concentration was applied to a 300-mesh Quantifoil R1.2/1.3 Cu grid pre-treated with glow-discharge, blotted in a Vitrobot Mark IV machine (force −5, time 3 s), and plunge-frozen in liquid ethane. The grid was loaded in a Titan Krios microscope equipped with Gatan BioQuantum K3 imaging filter and camera. A 20-eV slit was used for the filter. Data collection was done with serialEM (46). Images were recorded at 81,000× magnification, corresponding to a pixel size of 1.06 Å/pix. A defocus range of −1.0μm to −1.8μm was set. A total dose of 50 e-/Å2 of each exposure was fractionated into 50 frames. The first two frames of the movie stacks were not included in motion-correction. CryoEM data processing was performed on the fly with cryoSPARC Live (47) following regular single particle procedures.

### Virus propagation and titration

Methods from our previous publications were applied for DBV propagation and titration (16, 17). Briefly, Vero E6 (ATCC, CRL-1586) cells were cultured at 37°C and 5% CO_2_ with Dulbecco’s modified Eagle’s medium (DMEM) supplemented with 10% FBS. Cells were infected with CB1/2014 strain of DBV at confluency and supernatant was collected 7days later. The supernatant was centrifuged to remove cell debris and stored at −80°C until further use. Viral titer in form of TCID_50_ was determined by immunofluorescence assay (IFA) using in house-generated mouse monoclonal antibody recognizing DBV nucleoprotein (Np).

### Animal Care

BALB/c mice at ages of 6-8 weeks (Jackson Laboratories, Maine) were housed in Biological Resource Unit facility within Lerner Research Institute, Cleveland Clinic, with 12h light/dark cycle with access to water and diet.

Aged ferrets at ages of 48-50 months (ID Bio, Cheongju, South Korea) were housed in Laboratory Animal Research Center of Chungbuk National University (LARC) (Cheongju, South Korea) with 12h light/dark cycle with access to water and diet. All mouse and ferret cares were performed in accordance with the institutional animal care guideline and experiment protocols approved by Institutional Biosafety Committee (IBC) and Institutional Animal Care and Use Committee (IACUC) in Cleveland Clinic and Chungbuk National University, respectively. After viral challenge, the animals were monitored more frequently by the authors or veterinary technicians on duty. Viruses were handled in an enhanced biosafety level 3 containment laboratory as approved by the Korean Centers for Disease Control and Prevention (KCDC-14-3-07).

### Animal immunization and sample collection

Mice were intramuscularly immunized with 3.3μg of FT nanoparticles, or 1μg, 5μg, or 10μg of GnH-FT nanoparticles in hind leg. The antigens were prepared in 50μl of DPBS and mixed with 50μl AddaVax adjuvant (veterinary equivalent to MF59, Invivogen). Blood was collected from saphenous vein or retro orbital sinus to titer neutralizing antibodies.

For ferret immunization, 15μg FT nanoparticles or GnH-FT nanoparticles in 300μl was mixed with 300μl of AddaVax adjuvant for intramuscular immunization into the legs under anesthesia. Blood was also collected at the anesthesia. Subsequently, ferrets were intramuscularly infected with 10^7.6^ TCID_50_/mL of DBV, which has shown 100% fatality in our previous study (45). Their body weight and temperature were measured, and veterinary clinical symptoms were observed. Blood was collected for hematological analysis every other day until 14 days post infection. Three animals per group were sacrificed at days 2, 4, and 6 to collect serum, spleen, liver, and kidney with individual scissors to avoid cross-contamination.

### Titration of DBV Gn-recognizing antibodies and DBV-neutralizing antibodies in serum

To measure total mouse IgG against GnH, FT and GnH-FT, ELISA plates (MaxiSorp, ThermoFisher) were coated with respective antigens at concentration of 0.1μg/well. The plates were blocked with 5% skim milk in 0.05% PBS-Tween 20. Heat-inactivated sera was diluted in 10-fold dilution series in DPBS and 100μl of the dilutions were incubated in the wells overnight at 4°C. Plates were washed and incubated with HRP-conjugated anti-mouse IgG antibody (Jackson Immunoresearch). For detection of antibodies, the plates were overlaid with TMB substrate (ThermoFisher) and 1M sulfuric acid.

To measure mouse neutralizing antibody titer against DBV, we performed pseudovirus neutralization assay as described in our previous publication (40). In brief, we co-incubated serially 2-fold diluted sera with recombinant VSV carrying DBV glycoproteins and reporter luciferase gene (rVSV-DBV-Luc). The inoculum was added to HEK293T cells and incubated at 37 ° C with 5% CO_2_ and the luciferase signal was with analyzed using luciferase assay kit (Promega).

For titration of total ferret IgG and neutralizing antibodies against DBV, we performed ELISA and serum neutralization titration as previously described (16, 45). To titer total ferret IgG titer, ELISA plates coated with antigen and blocked in same method as from mouse IgG titer. Ferret sera were diluted in 2% skim milk in 0.05% PBS-Tween 20 from 1:50 to 1:50,000. 100μl of diluted ferret sera were incubated in ELISA plates for 2h at room temperature. Plates were then washed and incubated with HRP-conjugated anti-ferret IgG (KPL, South Korea). O-phenylenediamine dihydrochloride substrate was added to develop color and 1M sulfuric acid stop solution was added. After washing ELISA plates coated with GnH, the plates were blocked and incubated with HRP-conjugated anti-ferret IgG (KPL, South Korea). O-phenylenediamine dihydrochloride (ThermoFisher) was added to the plates and 1M sulfuric acid was added to stop color development. OD values at 450nm were measured with a plate reader (iMark Microplate reader, Bio-Rad).

Serum neutralization titer was measured as previously described (48). Briefly, heat-inactivated serum samples were serially twofold diluted from a 1:2 to 1:128. Then, 50μl of the diluted serum were mixed with equal volume of 200 focus-forming units of DBV for 1h at 37°C. The mixture was adsorbed onto confluent Vero E6 cells in 96-well plate at 37°C for 1h. Media was changed to maintenance medium and cells were maintained in incubator for 5 days. Cells were then fixed with 10% formalin and stained with in house-generated anti-DBV Np antibody. For FRNT_50_, the cells were also stained with HRP-conjugated anti-mouse IgG antibody. Serum neutralizing antibody titer was presented as reciprocal of the highest serum dilution neutralizing fluorescence signal of DBV Np.

### Profiling T-cell immunity by IFN-γ ELISpot

We referred to our previous publication (16). In brief, Multiscreen 96-well plates with PVDF membrane (Milipore) were coated with 100ul of anti-mouse IFN-γ antibody (clone AN-18, eBioscience, South Korea) overnight at 4°C. Mouse splenocytes were stimulated *ex vivo* with overlapping peptide pool (OLP) of 78 of 15-mer peptides covering DBV GnH formulated at 0.625μg/ml for each peptide in RPMI medium (Gibco) in the 96-well plate. PMA at 10ng/ml and ionomycin at 500ng/ml were included as positive controls and 0.5% DMSO was included as negative control for stimulation. After 24h stimulation in 5% CO_2_, 37°C incubator, plates were washed to remove cells and incubated with 100μl of biotinylated anti-mouse IFN-γ antibody for 1h at RT, followed by wash and incubation with 100μL of streptavidin-alkaline phosphatase (Invitrogen) for 1h at RT. 100ul of BCIP/NBT was added for 10min incubation at RT. Number of spot forming units (SFUs) per cells were presented after subtracting SFUs from negative control wells stimulated with 0.5% final concentration of DMSO.

### Intracellular cytokine staining

Based our protocol from previous publication (16), we resuspended the splenocytes from the immunized in 100μl of RPMI-1640 media. The cells were *ex vivo*-stimulated with anti-CD107a antibody (BD Biosciences, 553792), anti-CD28/CD49d antibody (BD Biosciences, 347690) and OLP or DMSO in 100μl. The mixture was incubated in 5% CO_2_, 37°C incubator for 1hour and treated with 4μl of mixture of the complete RPMI-1640 media:Brefeldin A (GolgiPlug, BD Biosciences, 555029):Monensin (GolgiStop, BD Biosciences, 554715) in 55:3:2. After 12hours of incubation in 5% CO_2_, 37°C, the cells were washed with PBS and stained with surface antibodies (anti-CD44 BV421 [BD Biosciences, 536970], anti-CD8a BV510 [BD Biosciences, 563068], anti-CD62L BV650 [BD Biosciences, 564108], anti-CD3 BV786 [BD Biosciences, 564010], anti-CD4 PerCP-Cy5.5 [BD Biosciences, 561115], and anti-CD19 APC [BD Biosciences, 561738]) for 15 minutes at RT and washed. Cells were then fixed with 4% paraformaldehyde in PBS and permeabilized. We then washed the cells and stained with FACS antibodies (anti-IL-2 FITC [BD Biosciences, 562040], anti-TNF PE [BD Biosciences, 554419], anti-IFN-γ [BD Biosciences, 557735], and anti-CD107a [BD Biosciences, 560647]) by 20 minutes of incubation at room temperature, washed twice with the permeabilization buffer and resuspended with 300μl of PBS.

### Hematological analysis and viral titration from challenged ferrets

We analyzed for hematological profile and tittered viremia as previously described (17). Total white blood cell and platelet counts in whole ferret blood samples were analyzed using the Celltac hematology analyzer (MEK-6550J/K, Nihon Kohden, Japan). Total RNA was extracted with TRIzol reagent (ThermoFisher) and reverse-transcribed to generate cDNA using QuantiTect Reverse Transcription system (Qiagen). Primers for real-time RT PCR (F: AATTCACATTTGAGGGTAGTT), R: TATCCAAGGAGGATGACAATAAT) were designed to recognize M segment of DBV genome. Real-time PCR was performed with SYBR Green supermix and CFX Real-Time PCR detection system (Bio-Rad). Copy numbers were normalized to GAPDH gene.

## Acknowledgments

*H. pylori*-bullfrog ferritin construct was kindly provided by Drs. Jeffrey Cohen and Gary Nabel at Vaccine Research Center, NIAID. This work was supported by National research foundation of Korea (NRF-2020R1A5A2017476, 2020R1A2C3008339), Institute for Basic Science (IBS) of Korea (IBS-R801-D1 to Y.K.C.), and National Institute of Health (CA251275, AI140705, AI140718, AI152190, AI171201, AI172252, DE023926, and DE028521) and Korea Research Institute of Bioscience and Biotechnology KGM9942011 (J.U.J.). We appreciate Carolyn Marks (Core Center of Excellence in Nano Imaging, University of Southern California) for technical assistance with transmission electron microscopy of nanoparticles.

---

**Fig S1. Structural analysis of FT and GnH-FT nanoparticles**

(A, B) Negatively stained scanning electron microscopy (EM) of FT (A) and GnH-FT (B) nanoparticles. Average diameters of the nanoparticles were calculated from 50 randomly selected nanoparticles’ diameters.

**Fig S2. Antibody response against FT and GnH-FT as antigens in immunized mice**

(A, B) Experiment from Fig 3B was repeated using FT (A) and GnH-FT (B) as antigens to test for total IgG response against FT and GnH-FT. Total IgG against FT stands for off-target antibody response which targets FT, instead of GnH, as the primary target of the immune response. The asterisks represent statistical significance of endpoint titer from mice immunized with GnH-FT-nanoparticle to mice immunized with FT evaluated with one-way ANOVA with Dunnett multiple comparison test.

*P < 0.05, **P < 0.01, ***P < 0.001 and ****P < 0.00001.

## References

1. Casel MA, Park SJ, Choi YK. 2021. Severe fever with thrombocytopenia syndrome virus: emerging novel phlebovirus and their control strategy. Experimental & Molecular Medicine 53:713–722.

2. Yu XJ, Liang MF, Zhang SY, Liu Y, Li JD, Sun YL, Zhang L, Zhang QF, Popov VL, Li C, Qu J, Li Q, Zhang YP, Hai R, Wu W, Wang Q, Zhan FX, Wang XJ, Kan B, Wang SW, Wan KL, Jing HQ, Lu JX, Yin WW, Zhou H, Guan XH, Liu JF, Bi ZQ, Liu GH, Ren J, Wang H, Zhao Z, Song JD, He JR, Wan T, Zhang JS, Fu XP, Sun LN, Dong XP, Feng ZJ, Yang WZ, Hong T, Zhang Y, Walker DH, Wang Y, Li DX. 2011. Fever with thrombocytopenia associated with a novel bunyavirus in China. N Engl J Med 364:1523–32.

3. Rattanakomol P, Khongwichit S, Linsuwanon P, Lee KH, Vongpunsawad S, Poovorawan Y. 2022. Severe Fever with Thrombocytopenia Syndrome Virus Infection, Thailand, 2019-2020. Emerg Infect Dis 28:2572–2574.

4. Tran XC, Yun Y, Van An L, Kim SH, Thao NTP, Man PKC, Yoo JR, Heo ST, Cho NH, Lee KH. 2019. Endemic Severe Fever with Thrombocytopenia Syndrome, Vietnam. Emerg Infect Dis 25:1029–1031.

5. Jiang XL, Zhang S, Jiang M, Bi ZQ, Liang MF, Ding SJ, Wang SW, Liu JY, Zhou SQ, Zhang XM, Li DX, Xu AQ. 2015. A cluster of person-to-person transmission cases caused by SFTS virus in Penglai, China. Clin Microbiol Infect 21:274–9.

6. Zhang YZ, He YW, Dai YA, Xiong Y, Zheng H, Zhou DJ, Li J, Sun Q, Luo XL, Cheng YL, Qin XC, Tian JH, Chen XP, Yu B, Jin D, Guo WP, Li W, Wang W, Peng JS, Zhang GB, Zhang S, Chen XM, Wang Y, Li MH, Li Z, Lu S, Ye C, de Jong MD, Xu J. 2012. Hemorrhagic fever caused by a novel Bunyavirus in China: pathogenesis and correlates of fatal outcome. Clin Infect Dis 54:527–33.

7. Yun SM, Park SJ, Kim YI, Park SW, Yu MA, Kwon HI, Kim EH, Yu KM, Jeong HW, Ryou J, Lee WJ, Jee Y, Lee JY, Choi YK. 2020. Genetic and pathogenic diversity of severe fever with thrombocytopenia syndrome virus (SFTSV) in South Korea. JCI Insight 5.

8. Li J, Li S, Yang L, Cao P, Lu J. 2021. Severe fever with thrombocytopenia syndrome virus: a highly lethal bunyavirus. Crit Rev Microbiol 47:112–125.

9. Anonymous. 2018. Annual review of diseases prioritized under the Research and Development Blueprint, Informal consultation. World Health Organization, https://www.who.int/docs/default-source/blue-print/2018-annual-review-of-diseases-prioritized-under-the-research-and-development-blueprint.pdf?sfvrsn=4c22e36_2.

10. Diseases NIoAaI. 2018. NIAID Emerging Infectious Diseases/Pathogens - Category C. Diseases BaEI,

11. Plegge T, Hofmann-Winkler H, Spiegel M, Pöhlmann S. 2016. Evidence that Processing of the Severe Fever with Thrombocytopenia Syndrome Virus Gn/Gc Polyprotein Is Critical for Viral Infectivity and Requires an Internal Gc Signal Peptide. PLoS One 11:e0166013.

12. Halldorsson S, Behrens AJ, Harlos K, Huiskonen JT, Elliott RM, Crispin M, Brennan B, Bowden TA. 2016. Structure of a phleboviral envelope glycoprotein reveals a consolidated model of membrane fusion. Proc Natl Acad Sci U S A 113:7154–9.

13. Guo X, Zhang L, Zhang W, Chi Y, Zeng X, Li X, Qi X, Jin Q, Zhang X, Huang M, Wang H, Chen Y, Bao C, Hu J, Liang S, Bao L, Wu T, Zhou M, Jiao Y. 2013. Human antibody neutralizes severe Fever with thrombocytopenia syndrome virus, an emerging hemorrhagic Fever virus. Clin Vaccine Immunol 20:1426–32.

14. Kim KH, Kim J, Ko M, Chun JY, Kim H, Kim S, Min JY, Park WB, Oh MD, Chung J. 2019. An anti-Gn glycoprotein antibody from a convalescent patient potently inhibits the infection of severe fever with thrombocytopenia syndrome virus. PLoS Pathog 15:e1007375.

15. Wu Y, Zhu Y, Gao F, Jiao Y, Oladejo BO, Chai Y, Bi Y, Lu S, Dong M, Zhang C, Huang G, Wong G, Li N, Zhang Y, Li Y, Feng WH, Shi Y, Liang M, Zhang R, Qi J, Gao GF. 2017. Structures of phlebovirus glycoprotein Gn and identification of a neutralizing antibody epitope. Proc Natl Acad Sci U S A 114:E7564–E7573.

16. Kwak J-E, Kim Y-I, Park S-J, Yu M-A, Kwon H-I, Eo S, Kim T-S, Seok J, Choi W-S, Jeong JH, Lee H, Cho Y, Kwon JA, Jeong M, Maslow JN, Kim Y-E, Jeon H, Kim KK, Shin E-C, Song M-S, Jung JU, Choi YK, Park S-H. 2019. Development of a SFTSV DNA vaccine that confers complete protection against lethal infection in ferrets. Nature Communications 10:3836.

17. Park SJ, Kim YI, Park A, Kwon HI, Kim EH, Si YJ, Song MS, Lee CH, Jung K, Shin WJ, Zeng J, Choi Y, Jung JU, Choi YK. 2019. Ferret animal model of severe fever with thrombocytopenia syndrome phlebovirus for human lethal infection and pathogenesis. Nat Microbiol 4:438–446.

18. Choi Y, Jiang Z, Shin WJ, Jung JU. 2020. Severe Fever with Thrombocytopenia Syndrome Virus NSs Interacts with TRIM21 To Activate the p62-Keap1-Nrf2 Pathway. J Virol 94.

19. Zimmermann P, Curtis N. 2019. Factors That Influence the Immune Response to Vaccination. Clinical Microbiology Reviews 32:e00084–18.

20. Pollard AJ, Bijker EM. 2021. A guide to vaccinology: from basic principles to new developments. Nature Reviews Immunology 21:83–100.

21. Gause KT, Wheatley AK, Cui J, Yan Y, Kent SJ, Caruso F. 2017. Immunological Principles Guiding the Rational Design of Particles for Vaccine Delivery. ACS Nano 11:54–68.

22. Porter CJH, Trevaskis NL. 2020. Targeting immune cells within lymph nodes. Nature Nanotechnology 15:423–425.

23. Kelly HG, Tan HX, Juno JA, Esterbauer R, Ju Y, Jiang W, Wimmer VC, Duckworth BC, Groom JR, Caruso F, Kanekiyo M, Kent SJ, Wheatley AK. 2020. Self-assembling influenza nanoparticle vaccines drive extended germinal center activity and memory B cell maturation. JCI Insight 5.

24. Link A, Zabel F, Schnetzler Y, Titz A, Brombacher F, Bachmann MF. 2012. Innate immunity mediates follicular transport of particulate but not soluble protein antigen. J Immunol 188:3724–33.

25. Kim YI, Kim D, Yu KM, Seo HD, Lee SA, Casel MAB, Jang SG, Kim S, Jung W, Lai CJ, Choi YK, Jung JU. 2021. Development of Spike Receptor-Binding Domain Nanoparticles as a Vaccine Candidate against SARS-CoV-2 Infection in Ferrets. mBio 12.

26. Kanekiyo M, Bu W, Joyce MG, Meng G, Whittle JR, Baxa U, Yamamoto T, Narpala S, Todd JP, Rao SS, McDermott AB, Koup RA, Rossmann MG, Mascola JR, Graham BS, Cohen JI, Nabel GJ. 2015. Rational Design of an Epstein-Barr Virus Vaccine Targeting the Receptor-Binding Site. Cell 162:1090–100.

27. Kanekiyo M, Wei CJ, Yassine HM, McTamney PM, Boyington JC, Whittle JR, Rao SS, Kong WP, Wang L, Nabel GJ. 2013. Self-assembling influenza nanoparticle vaccines elicit broadly neutralizing H1N1 antibodies. Nature 499:102–6.

28. Kim YS, Son A, Kim J, Kwon SB, Kim MH, Kim P, Kim J, Byun YH, Sung J, Lee J, Yu JE, Park C, Kim YS, Cho NH, Chang J, Seong BL. 2018. Chaperna-Mediated Assembly of Ferritin-Based Middle East Respiratory Syndrome-Coronavirus Nanoparticles. Front Immunol 9:1093.

29. Domnich A, Arata L, Amicizia D, Puig-Barbera J, Gasparini R, Panatto D. 2017. Effectiveness of MF59-adjuvanted seasonal influenza vaccine in the elderly: A systematic review and meta-analysis. Vaccine 35:513–520.

30. Podda A, Del Giudice G. 2003. MF59-adjuvanted vaccines: increased immunogenicity with an optimal safety profile. Expert Review of Vaccines 2:197–204.

31. Pellegrini M, Nicolay U, Lindert K, Groth N, Della Cioppa G. 2009. MF59-adjuvanted versus non-adjuvanted influenza vaccines: integrated analysis from a large safety database. Vaccine 27:6959–65.

32. Tsai TF. 2013. Fluad(R)-MF59(R)-Adjuvanted Influenza Vaccine in Older Adults. Infect Chemother 45:159–74.

33. Maini MK, Pallett LJ. 2018. Defective T-cell immunity in hepatitis B virus infection: why therapeutic vaccination needs a helping hand. Lancet Gastroenterol Hepatol 3:192–202.

34. Thimme R. 2021. T cell immunity to hepatitis C virus: Lessons for a prophylactic vaccine. J Hepatol 74:220–229.

35. (CDC) CfDCaP. 2023. What you need to know about Asian longhorned ticks - a new tick in the United States.

36. Zhang X, Zhao C, Cheng C, Zhang G, Yu T, Lawrence K, Li H, Sun J, Yang Z, Ye L, Chu H, Wang Y, Han X, Jia Y, Fan S, Kanuka H, Tanaka T, Jenkins C, Gedye K, Chandra S, Price DC, Liu Q, Choi YK, Zhan X, Zhang Z, Zheng A. 2022. Rapid Spread of Severe Fever with Thrombocytopenia Syndrome Virus by Parthenogenetic Asian Longhorned Ticks. Emerg Infect Dis 28:363–372.

37. Yun SM, Lee WG, Ryou J, Yang SC, Park SW, Roh JY, Lee YJ, Park C, Han MG. 2014. Severe fever with thrombocytopenia syndrome virus in ticks collected from humans, South Korea, 2013. Emerg Infect Dis 20:1358–61.

38. Kimura M, Egawa K, Ozawa T, Kishi H, Shimojima M, Taniguchi S, Fukushi S, Fujii H, Yamada H, Tan L, Sano K, Katano H, Suzuki T, Morikawa S, Saijo M, Tani H. 2021. Characterization of pseudotyped vesicular stomatitis virus bearing the heartland virus envelope glycoprotein. Virology 556:124–132.

39. Lam TT, Liu W, Bowden TA, Cui N, Zhuang L, Liu K, Zhang YY, Cao WC, Pybus OG. 2013. Evolutionary and molecular analysis of the emergent severe fever with thrombocytopenia syndrome virus. Epidemics 5:1–10.

40. Xia T, Wu X, Hong E, Jung K, Lai C-J, Kwak M-J, Seo H, Kim S, Jiang Z, Cha I, Jung JU. 2023. Glucosylceramide is essential for Heartland and Dabie bandavirus glycoprotein-induced membrane fusion. PLOS Pathogens 19:e1011232.

41. Yoshikawa T, Shimojima M, Fukushi S, Tani H, Fukuma A, Taniguchi S, Singh H, Suda Y, Shirabe K, Toda S, Shimazu Y, Nomachi T, Gokuden M, Morimitsu T, Ando K, Yoshikawa A, Kan M, Uramoto M, Osako H, Kida K, Takimoto H, Kitamoto H, Terasoma F, Honda A, Maeda K, Takahashi T, Yamagishi T, Oishi K, Morikawa S, Saijo M. 2015. Phylogenetic and Geographic Relationships of Severe Fever With Thrombocytopenia Syndrome Virus in China, South Korea, and Japan. J Infect Dis 212:889–98.

42. Li A, Liu L, Wu W, Liu Y, Huang X, Li C, Liu D, Li J, Wang S, Li D, Liang M. 2021. Molecular evolution and genetic diversity analysis of SFTS virus based on next-generation sequencing. Biosafety and Health 3:105–115.

43. Yun SM, Park SJ, Park SW, Choi W, Jeong HW, Choi YK, Lee WJ. 2017. Molecular genomic characterization of tick- and human-derived severe fever with thrombocytopenia syndrome virus isolates from South Korea. PLoS Negl Trop Dis 11:e0005893.

44. Jia Z, Wu X, Wang L, Li X, Dai X, Liang M, Cao S, Kong Y, Liu J, Li Y, Wang J. 2017. Identification of a candidate standard strain of severe fever with thrombocytopenia syndrome virus for vaccine quality control in China using a cross-neutralization assay. Biologicals 46:92–98.

45. Kwang-Min Yu S-JP, Min-Ah Yu, Young-Il Kim, Younho Choi, Jae U. Jung, Benjamin Brennan, Young Ki Choi. 2019. Cross-genotype protection of live-attenuated vaccine candidate for severe fever with thrombocytopenia syndrome virus in a ferret model. PNAS 116:26900–26908.

46. Mastronarde DN. 2005. Automated electron microscope tomography using robust prediction of specimen movements. J Struct Biol 152:36–51.

47. Punjani A, Rubinstein JL, Fleet DJ, Brubaker MA. 2017. cryoSPARC: algorithms for rapid unsupervised cryo-EM structure determination. Nat Methods 14:290–296.

48. Yu KM, Yu MA, Park SJ, Kim YI, Robles NJ, Kwon HI, Kim EH, Si YJ, Nguyen HD, Choi YK. 2018. Seroprevalence and genetic characterization of severe fever with thrombocytopenia syndrome virus in domestic goats in South Korea. Ticks Tick Borne Dis 9:1202–1206.

