## Supplementary figures 1 and 2 for "Self-assembling Gn head ferritin nanoparticle vaccine provides full protection from lethal challenge of Dabie Bandavirus in aged ferrets"

### Slide 1
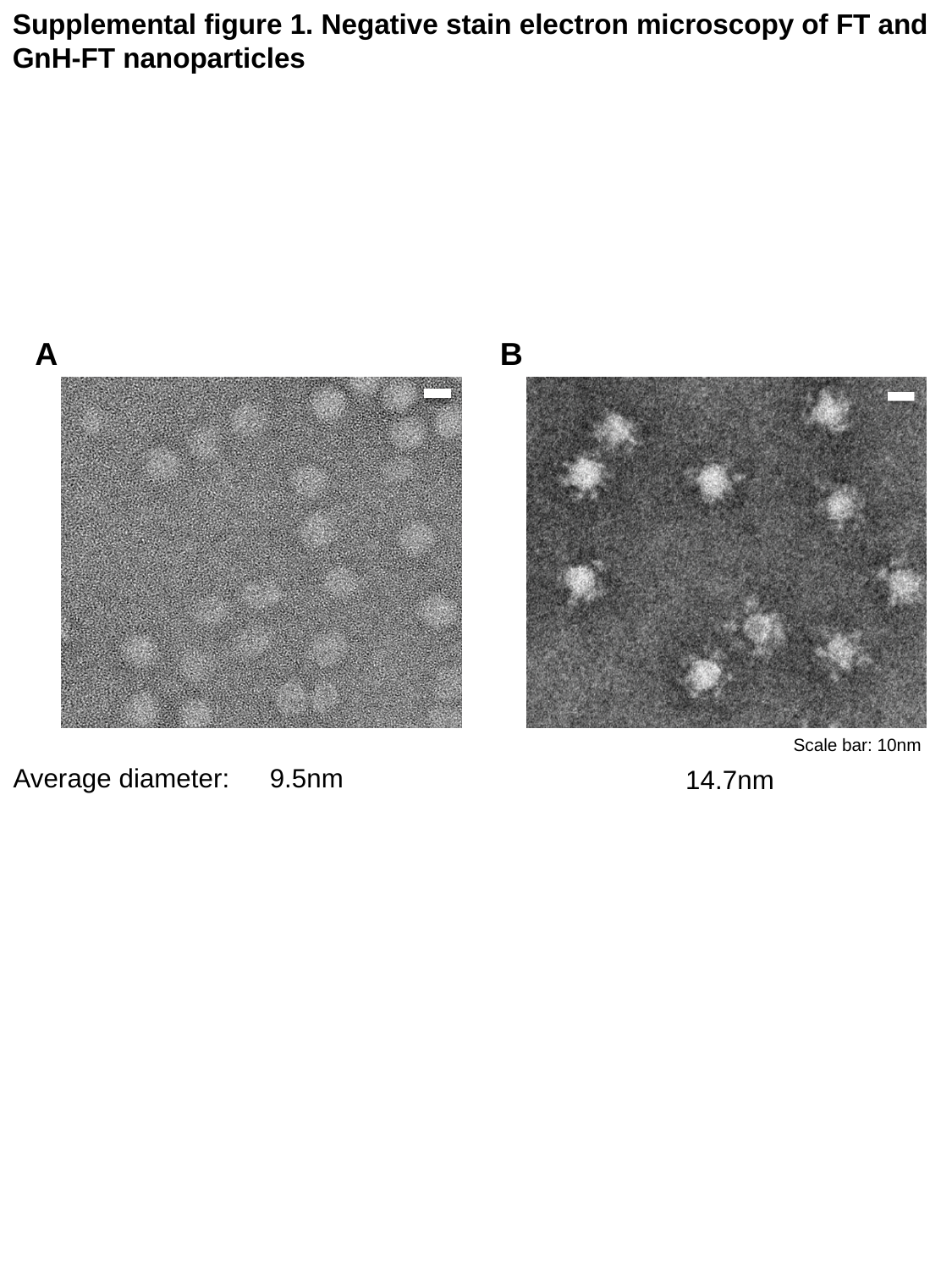

Supplemental figure 1. Negative stain electron microscopy of FT and GnH-FT nanoparticles
A
B
Scale bar: 10nm
Average diameter:
9.5nm
14.7nm

### Slide 2
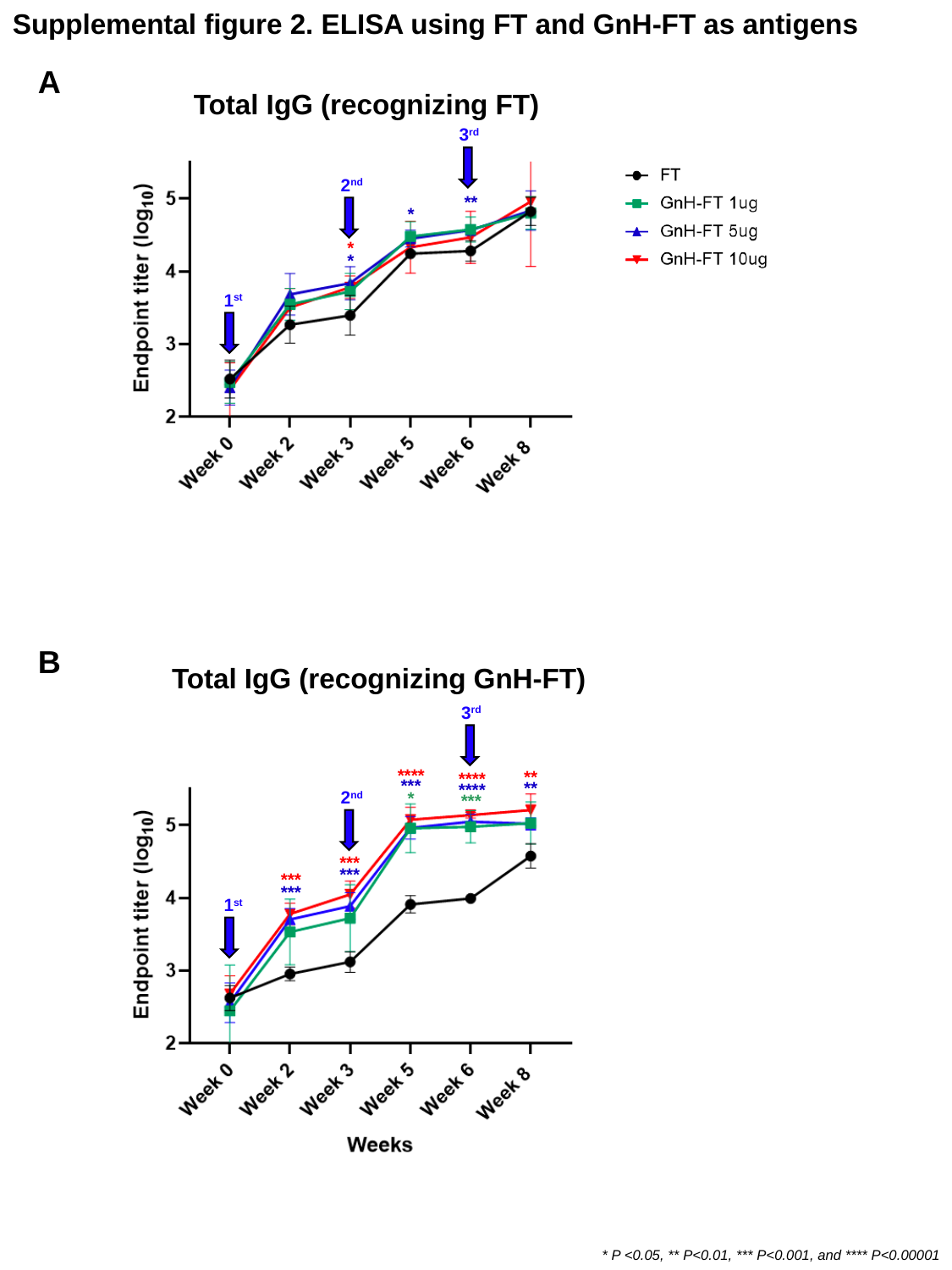

Supplemental figure 2. ELISA using FT and GnH-FT as antigens
A
Total IgG (recognizing FT)
3^rd^
2^nd^
**
*
*
*
1^st^
B
Total IgG (recognizing GnH-FT)
3^rd^
****
**
****
***
**
****
2^nd^
*
***
***
***
***
***
1^st^
* P <0.05, ** P<0.01, *** P<0.001, and **** P<0.00001
